## Supplementary material for "Neonatal sensitivity to vocal emotions: A developmental change at 37 weeks of gestational age": data information

**Supplemental materials**

### Result of the three-way ANOVA

A three-way repeated measures ANOVA was conducted on the mean amplitudes of the MMR with condition (vocal/nonvocal) and hemisphere (left/right frontal, i.e., F3/F4) as within-subjects factors and neonatal group (GA = 35, 36, 37, 38, 39, and 40 weeks) as the between-subjects factor. The main effect of hemisphere is not significant, F(1,114) = 0.153, p = 0.696, $\eta_{p}^{2}$ = 0.001. The interaction between hemisphere and group is not significant, F(5,114) = 0.249, p = 0.940, $\eta_{p}^{2}$ = 0.011. The interaction between of hemisphere and stimuli is not significant, F(1,114) = 0.474, p = 0.492, $\eta_{p}^{2}$ = 0.004. The three-way interaction is not significant, F(5,114) = 0.666, p = 0.650, $\eta_{p}^{2}$ = 0.028.

### Epoch number

Epoch numbers in different conditions are reported in Table S1. For happy trials, a six (group) × two (condition: vocal/nonvocal) ANOVA was performed. Neither the main effect of condition (F(1,114) = 0.533, *p* = 0.467, $\eta_{p}^{2}$ = 0.005) nor group (F(5,114) = 0.795, *p* = 0.555, $\eta_{p}^{2}$ = 0.034) was found to be significant. The interaction between group and condition was also not significant (F(5,114) = 1.654, *p* = 0.151, $\eta_{p}^{2}$ = 0.068). For neutral trials, another 6 × 2 ANOVA was performed. Again, neither the main effect of condition (F(1,114) =1.137, *p* = 0.289, $\eta_{p}^{2}$ = 0.010) nor group (F(5,114) = 1.225, *p* = 0.302, $\eta_{p}^{2}$ = 0.051) was significant. The interaction between group and condition was not significant (F(5,114) = 1.012, *p* = 0.414, $\eta_{p}^{2}$ = 0.043).

Table S1. Epoch numbers in different conditions (mean ± standard deviation)

| GA group  (week) | vocal  happy | nonvocal  happy | vocal  neutral | nonvocal  neutral |
| --- | --- | --- | --- | --- |
| 35 | 53.15 ± 4.86 | 52.65 ± 6.79 | 215.55 ± 16.58 | 214.10 ± 21.80 |
| 36 | 51.20 ± 7.18 | 48.80 ± 11.04 | 201.90 ± 30.78 | 196.25 ± 38.24 |
| 37 | 48.45 ± 6.79 | 50.40 ± 5.98 | 191.40 ± 38.00 | 199.70 ± 32.30 |
| 38 | 48.35 ± 9.03 | 51.00 ± 6.47 | 197.95 ± 35.37 | 202.60 ± 27.40 |
| 39 | 49.40 ± 7.06 | 49.65 ± 8.11 | 198.35 ± 31.65 | 198.25 ± 36.47 |
| 40 | 49.80 ± 6.53 | 50.35 ± 5.71 | 192.95 ± 32.55 | 202.85 ± 24.42 |
